# Controlled expression of avian migratory fattening influences innate immune responses

**DOI:** 10.1101/2023.05.15.540777

**Authors:** Marcin Tobolka, Zuzanna Zielińska, Leonida Fusani, Nikolaus Huber, Ivan Maggini, Gianni Pola, Valeria Marasco

## Abstract

While immunity is frequently suppressed when birds engage in strenuous migratory flights, whether and how immunity changes during the rapid accumulation of energy stores in preparation for migration remains largely unknown. Here, we induced pre-migratory fattening through controlled changes of daylight in common quails *Coturnix coturnix* and regularly assessed changes in a marker of constitutive innate immunity (Leukocyte Coping Capacity or LCC) and measures of body composition (lean and fat mass). LCC responses were highest in the mid-fattening phase and lowest when fattening was completed. At mid-fattening, we also found that the birds that kept a higher proportion of lean mass (i.e. accumulated less fat) had the highest LCC peaks. Our results indicate that migratory birds undergo rapid immunological changes as they accumulate energy stores for migration and propose that this could be due to competing or trade-off processes between metabolic remodelling and innate immune system function.

**Summary statement:** Immunity is vital when migrating to new environments. It is costly and competivite to other physiological processes. Here we bring new evidence on this process in migratory birds.

## 1. Introduction

Birds show various physiological adaptations to accommodate the intense energy demands of long-distance migratory flights (Cornelius et al., 2013). One of the most spectacular preparatory features described so far is pre-migratory fattening – a rapid accumulation of energy stores prior to the onset of active migration (Bairlein, 2002; Jenni and Jenni-Eiermann, 1998; Jenni and Schaub, 2003). Studies across different bird species sampled at stopover sites (i.e. where migrating birds temporarily suspend the long-distance endurance flight to rest and refuel) show that individuals with larger fat stores can migrate faster (Brust et al., 2022; Sjöberg et al., 2015). Thus, the physiological and metabolic preparations (e.g. hyperphagia, fattening, and flight muscle hypertrophy) are vital for successful migration (Bairlein, 2002; Guglielmo, 2018; Hedenström and Alerstam, 1997).

Due to their mobility, migratory animals are exposed to a greater number and variety of pathogens compared to non-migratory populations (Buehler et al., 2010; van Dijk et al., 2014). Thus, physiological adaptations associated with the maintenance of immune system functioning are particularly relevant to minimise negative effects of infectious diseases in migratory species (Eikenaar et al., 2023; Fritzsche McKay and Hoye, 2016; Roitt et al., 2001). However, maintaining a high immune response might require adjustments in the amount of resources allocated to the system relative to other competing behavioural and physiological processes (Germain, 2012; Norris and Evans, 2000; Sheldon and Verhulst, 1996). For example, nutritional deficiencies often alter the development of immune system and impair its optimal functioning (Hotamisligil, 2006; Lochmiller et al., 1993; Mauck et al., 2005; SamartIn and Chandra, 2001). Adequate fine-tuning of effective immune responses is likely to be even more difficult in migratory species because they have to rapidly adjust their energy requirements in preparation for migration and between repeated stopovers and strenuous migratory flights (Altizer et al., 2011; Buehler et al., 2010; Eikenaar et al., 2023; Srygley and Lorch, 2011; Voigt et al., 2020).

Several studies in birds suggested that migrants temporarily suppress immunity and re-allocate energy to endurance flights (Altizer et al., 2011; Eikenaar and Hegemann, 2016; Eikenaar et al., 2018; Møller et al., 2004; Nebel et al., 2012; Nebel et al., 2013; Owen and Moore, 2006; Owen and Moore, 2008a; Owen and Moore, 2008b). However, it is still not clear whether and how immunity might be optimally modulated during pre-migratory fuelling, and to which extent individual variation of innate immunity is related to the rapid accumulation of energy stores during this life cycle stage. Recent studies in the northern wheatear *Oenanthe oenanthe* showed that energy stores of free-living birds sampled at a stopover site during autumn migration were positively correlated with immunity, and *ad libitum* feeding of temporarily caged individuals led to rapid increases in their blood bacterial killing capacity (Eikenaar et al., 2020a; Eikenaar et al., 2020b). It could be hypothesized that the accumulation of energy stores is associated with a potentiation of immune responses. On the other hand, immunological variation during premigratory fuelling might also be the consequence of a potential competition between the energy required for fat accumulation and/or muscle volume increases and the need to maintain a competent immune system. Here, we explored these two possibilities by experimentally controlling the migratory state of captive young adult common quails *Coturnix coturnix* (Linnaeus, 1758) to simulate autumn pre-migratory fattening. We assessed potential changes in leukocyte oxidative burst capacity [Leucocyte Coping Capacity - hereafter LCC - a measure of constitutive innate immune protection in response to a secondary external stimulus (McLaren et al., 2003)] - by collecting blood samples at three different physiological states: (1) before premigratory fattening, (2) at mid-pre-migratory fattening, and (3) at the pre-migratory fattening peak. We assessed whether variation in LCC responses was associated with within-individual changes in body composition (lean and fat mass) over these three phases.

## 2. Materials and Methods

### Ethical Statement

The experiment was performed in compliance with the Austria legislation with approval of the Ethics Committee of the University of Veterinary Medicine Vienna and the Federal Ministry of Science, Research and Economy (2021-0.466.199).

### Study subjects and photoperiod manipulation

Fertile eggs of common quails were obtained from a breeding stock population kept at the Istituto Sperimentale Zootecnico per la Sicilia (ISZS, Palermo, Italy), which originated from wild common quail founders (Smith et al., 2018). After transportation to the Konrad Lorenz Institute of Ethology (Vetmeduni, Vienna), the eggs were artificially incubated and hatchlings reared under a 16:8 hrs light:dark cycle until four weeks of age, when they were sexed and moved to enclosures of 80 x 100 x 210 cm in sex-mixed groups of 11-12 birds until the termination of the experiment. Food (turkey starter, Lagerhaus, Austria) and water were always provided *ad libitum*. All birds were maintained in photoperiod- and climate-controlled indoor animal facility rooms at 20-24°C. When the birds were six weeks of age (i.e. week 0 of pre-migration fattening), they were exposed to a gradual reduction of day length (30 min/week) over eight consecutive weeks until the photoperiod reached 12:12 hrs light:dark. This light:dark schedule simulates autumn migration and triggers pre-migratory fattening (Marasco et al., 2021).

### Measurement of whole blood Leukocyte Coping Capacity (LCC)

LCC measurements were performed on a subset of un-manipulated control quails selected from a larger study investigating the effects of food manipulations on migratory fattening (to be published elsewhere). Birds were sampled up to three times for LCC measurements: at week 0 (n = 18, 8 females and 10 males) at 16:8 hrs light:dark, at week 4 (n = 16, 6 females and 10 males) at 14:10 hrs light:dark, and at week 8 (n = 18, 7 females and 11 males) at 12:12 hrs light:dark. Part of the birds sampled at week 0 (n = 10) were used for separate experiments and were excluded from analyses at week 4 and at week 8. The same birds were sampled between week 4 and week 8; at week 4, we did not sample two birds as they showed signs of intense pecking, which were resolved by brief social isolation and therefore they were included in the sampling at week 8. One day before blood sampling, the birds housed within the same enclosures were transferred into single cages (71 x 109 x 79 cm), but all birds remained in visual and acoustic contact with their conspecifics to minimise possible effects of the changes in the social environment. All birds were sampled within three minutes from opening the cage, after which we immediately performed LCC measurements in the freshly collected whole blood samples (Huber et al., 2017 - full details in Supplementary Material). We excluded three LCC measurements from the statistics (one bird at week 4, two birds at week 8) because the control sample yielded extreme, unrealistic chemiluminescence levels. From the resulting LCC response curve, we extracted the LCC peak, which was defined as the maximum ROS production in the course of the induced oxidative burst. This variable was a reliable proxy of the entire LCC response over time (adj. R^2^ = 0.94, *p* < 0.0001, see Fig S1 and S2 in Supplementary Material), as shown in previous work in birds and mammals (Huber et al., 2020; Huber et al., 2019).

### Measurements of body mass and body composition

After collecting the blood sample, each bird was weighed to the nearest 0.01g using an electronic balance. Subsequently, its lean and fat mass were measured to the nearest 0.01 g using a quantitative magnetic resonance analyser (Echo-MRI™ Birds and Bats Body Composition Analyzer, Houston, USA) (Guglielmo et al., 2011). As at week 8 the birds were immediately euthanized for tissue analyses for separate experiments, the Echo-MRI measurements at week 8 were performed 2-4 days before the birds were blood sampled (the body mass of the birds on the day of the Echo-MRI scans and at the time of blood sampling were highly correlated: R = 0.98, *p* < 0.0001). The birds were placed into appropriately sized ventilated holding tubes (62 mm I.D.) inside the analyzer for approximately 3 min; each bird was scanned three times, and the average of the measurements was used for statistical analyses.

### Statistical analysis

Statistical analyses were performed in R v 3.6.2 (R Core Team, 2022) in RStudio v 1.3.1093 (R Studio Team, 2022) using General Linear Models (GLMs) or Generalized Linear Mixed Models (GLMMs) with a Gaussian distribution error [package “lme4” - (Bates et al., 2015)]. We assessed the effects of the photoperiod manipulation on proxies of migratory fattening (i.e. body mass, fat mass, and lean mass as well as fat mass and lean mass as a proportion of body mass) and LCC peak levels using separate GLMMs. We entered time (week 0, week 4, and week 8), sex (male, female) and their interaction as fixed factors, while individual bird identity was entered as a random factor to control for the presence of repeated measurements. As the interaction of time and sex was always not significant (*p* > 0.05) it was subsequently removed from the final models. LCC peak levels were log-transformed to improve model residuals. To assess the strength of within-individual effects, the GLMM model for LCC analysis was re-run using only the subset of birds that were repeatedly sampled across the three sampling time points (8 birds in total). To further explore the effect of photoperiod on LCC peak levels found in these main models, we performed two separate GLMs to test whether LCC peak levels at week 4 or at week 8 were associated with the rate of fattening preceding the relevant blood sampling time (i.e. calculated as the change in the proportion of fat, or lean mass in relation to body mass). As the change in lean and fat mass (% of body mass) between week 0-week 4 and week 4-week 8 were highly correlated (-0.97 < R < -0.74, *p* < 0.0001), we tested each of these covariates separately.

## 3. Results

### Effects of photoperiod manipulation on pre-migratory fattening

Body and fat mass increased linearly in relation to shortening daylength in both male and female quails (Table 1a, c; Figure 1a, c). Regardless of sex, lean mass increased during the first phase of pre-migratory fattening (week 0-week 4), but remained relatively stable in the following four weeks, week 4-week 8 (Table 1b; Figure 1b). The proportion of lean mass to body mass linearly decreased over time (Table 1d; Figure 1d) and the proportion of fat mass to body mass showed the opposite pattern (Table 1e; Figure 1e).

**Table 1.** Results of Generalized Linear Mixed Model with a Gaussian error distribution to assess the effects of photoperiod manipulation on (a) body mass, (b) lean mass, (c) fat mass, (d) proportion of lean mass, (e) proportion of fat mass, and (f) LCC peak response in a captive population of common quails to simulate autumn migratory fattening. Fixed factor estimates are indicated in parenthesis, r indicates random factor (intercept) and its associated variance. Significant terms (*p* < 0.05) are in bold.

|  | **Estimate** | **SE** | **DF** | **t** | **p** |
| --- | --- | --- | --- | --- | --- |
| Ring identity (r) | 52.340 |  |  |  |  |
| Residual | 66.600 |  |  |  |  |
| Intercept | 78.754 | 3.303 | 38.117 | 23.840 | <0.0001 |
| **Photoperiod (week 4)** | 22.266 | 3.168 | 39.829 | 7.029 | <0.0001 |
| **Photoperiod (week 8)** | 37.479 | 3.100 | 41.023 | 12.091 | <0.0001 |
| Sex (male) | -3.43 | 3.729 | 28.516 | -0.92 | 0.365 |

(b) Lean mass
|  | **Estimate** | **SE** | **DF** | **t** | **p** |
| --- | --- | --- | --- | --- | --- |
| Ring identity (r) | 30.570 |  |  |  |  |
| Residual | 20.440 |  |  |  |  |
| Intercept | 72.522 | 2.182 | 33.796 | 33.242 | <0.0001 |
| **Photoperiod (week 4)** | 8.340 | 1.841 | 34.557 | 4.531 | <0.0001 |
| **Photoperiod (week 8)** | 9.651 | 1.810 | 35.963 | 5.331 | <0.0001 |
| Sex (male) | -2.080 | 2.550 | 25.702 | -0.816 | 0.422 |

(c) Fat mass
|  | **Estimate** | **SE** | **DF** | **t** | **p** |
| --- | --- | --- | --- | --- | --- |
| Ring identity (r) | 30.040 |  |  |  |  |
| Residual | 110.490 |  |  |  |  |
| Intercept | 0.794 | 3.503 | 40.624 | 0.227 | 0.822 |
| **Photoperiod (week 4)** | 11.592 | 3.83 | 42.615 | 3.027 | 0.004 |
| **Photoperiod (week 8)** | 27.158 | 3.727 | 43.43 | 7.287 | <0.0001 |
| Sex (male) | -1.456 | 3.743 | 28.285 | -0.389 | 0.700 |

(d) Lean mass (%)
|  | **Estimate** | **SE** | **DF** | **t** | **p** |
| --- | --- | --- | --- | --- | --- |
| Ring identity (r) | 0.003 |  |  |  |  |
| Residual | 0.006 |  |  |  |  |
| Intercept | 0.926 | 0.03 | 39.329 | 31.292 | <0.0001 |
| **Photoperiod (week 4)** | -0.117 | 0.03 | 41.234 | -3.904 | 0.0003 |
| **Photoperiod (week 8)** | -0.206 | 0.029 | 42.287 | -7.065 | <0.0001 |
| Sex (male) | 0.003 | 0.033 | 28.907 | 0.079 | 0.938 |

(e) Fat mass (%)
|  | **Estimate** | **SE** | **DF** | **t** | **p** |
| --- | --- | --- | --- | --- | --- |
| Ring identity (r) | 0.002 |  |  |  |  |
| Residual | 0.007 |  |  |  |  |
| Intercept | 0.006 | 0.038 | 40.463 | 0.230 | 0.819 |
| **Photoperiod (week 4)** | 0.111 | 0.031 | 42.498 | 3.627 | 0.001 |
| **Photoperiod (week 8)** | 0.223 | 0.030 | 43.324 | 7.518 | <0.0001 |
| Sex (male) | -0.010 | 0.030 | 27.871 | -0.344 | 0.733 |

(f) LCC peak
|  | **Estimate** | **SE** | **DF** | **t** | **p** |
| --- | --- | --- | --- | --- | --- |
| Ring identity (r) | 0.715 |  |  |  |  |
| Residual | 0.554 |  |  |  |  |
| Intercept | 4.659 | 0.346 | 35.697 | 13.451 | <0.0001 |
| **Photoperiod (week 4)** | 0.754 | 0.304 | 33.839 | 2.483 | 0.018 |
| **Photoperiod (week 8)** | -0.741 | 0.306 | 35.580 | -2.422 | 0.021 |
| Sex (male) | 0.509 | 0.403 | 28.607 | 1.263 | 0.217 |

**Figure 1.**
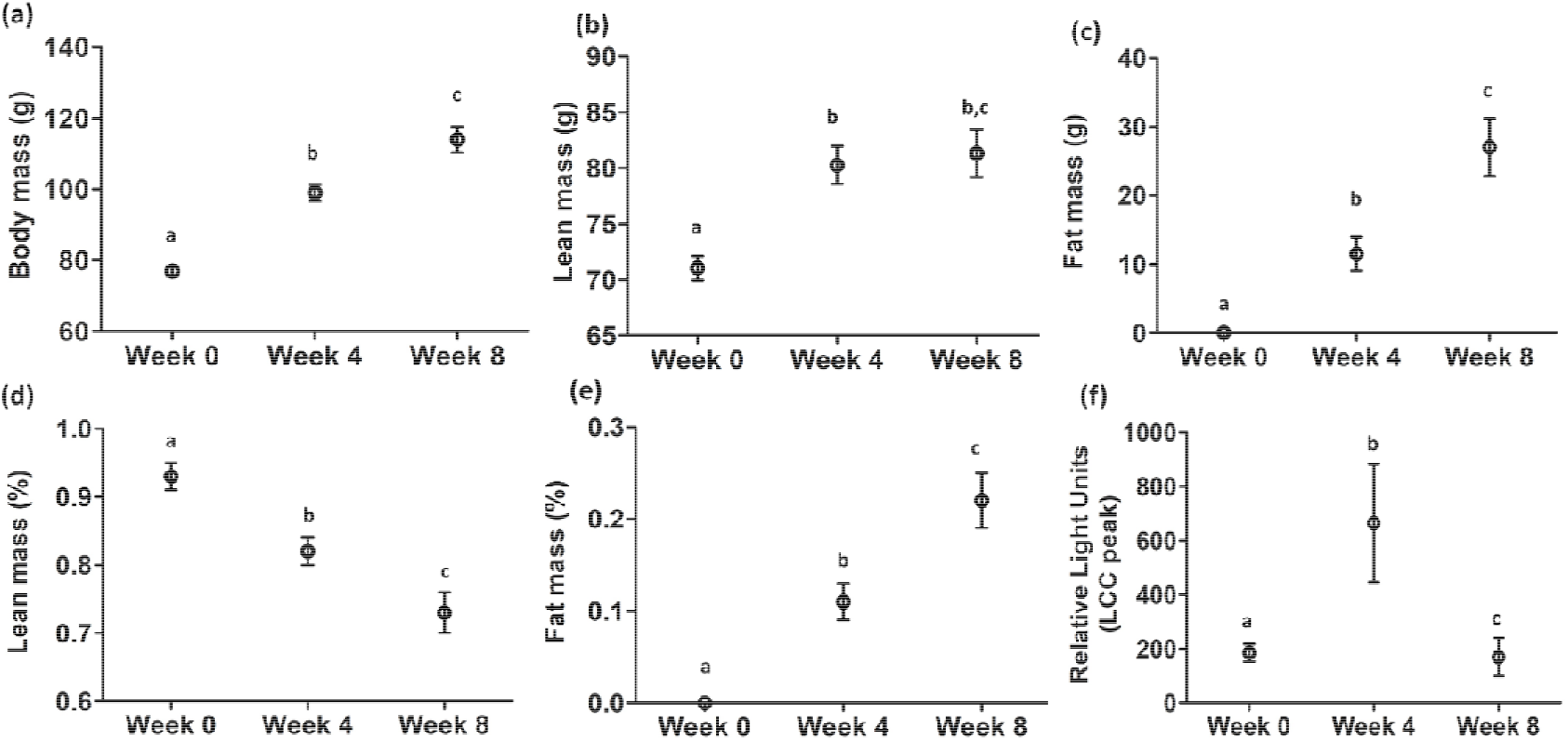
Overall body mass, lean and fat masses, and LCC peaks varied significantly over the experimental stages. Changes in (a) body mass (g), (b) lean mass (g), (c) fat mass (g), (d) lean mass as % of body mass, (e) fat mass as % of body mass, and LCC peak levels at each sampling time point, before the start of pre-migratory fattening (week 0), at mid-pre-migratory fattening (week 4), and at the pre-migratory fattening peak (week 8). Sample sizes in (a)-(e): week 0, n = 18; week 4, n = 16, and week 8, n = 18; in (f): week 0, n = 18, week 4, n = 15, and week 8, n = 16. Different letters indicate pairwise statistical differences (*p* < 0.05).

### Effects of photoperiod manipulation on LCC peak levels

Irrespective of sex, LCC peak levels differed among sampling weeks (Table 1f). LCC peak values were on average higher at week 4 compared to week 0 (*p* = 0.02, Figure 1f) and they were lower at week 8 compared to week 4 (*p <* 0.0001, Figure 1f) as well as week 0 (*p* = 0.02, Figure 1f). We obtained similar model estimates when using only the subset of birds that were repeatedly sampled across the three sampling time points (Table S1, Supplementary Material Material).

Our post-hoc analysis revealed that the relatively larger variation of LCC peak levels found at week 4 was associated with the change in lean mass relative to body mass since the start of the fattening process (estimate ± SE: 4733.1±1869.3, t = 2.532, *p* = 0.03; Figure 2a). The birds with higher LCC peak responses showed lower decreases in lean mass proportion between week 0 and week 4. We found a negative correlation between LCC peak responses measured at week 4 and the change in fat mass (as % of body mass) between week 0 and week 4. Birds with higher LCC peaks showed lower proportional increases in fat mass, though this pattern was not statistically significant (estimate ± SE: -4707±2292.5, t = -2.1, *p* = 0.06; Figure 2b). LCC peak levels at week 8 were not related to either change in lean mass proportion (estimate ± SE: 280.8±562.0, t = 0.5, *p* = 0.6; Figure 2c) or fat mass proportion (estimate ± SE: -508.6±528.0, t = -0.96, *p* = 0.4; Figure 2d) observed between week 4 - week 8.

**Figure 2.**
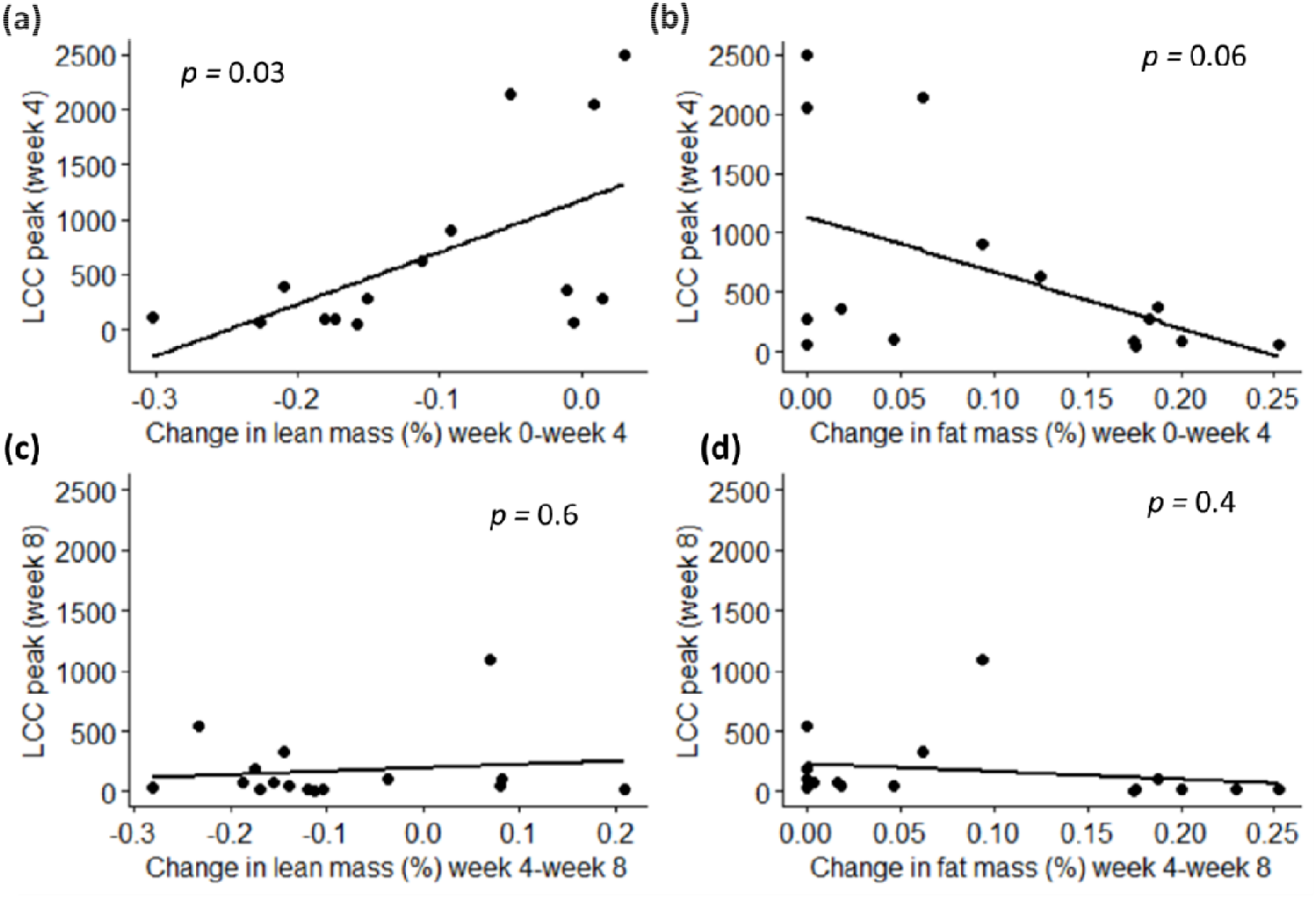
LCC peak was positively correlated with change in lean mass proportion, and negatively with fat mass proportion at the week 4 of the experiment. Correlation plots between LCC peak responses observed at week 4 (a-b) and at week 8 (c-d) and changes in lean mass as proportion of body mass (lean mass %) or fat mass as proportion of body mass (fat mass %) in the periods preceding the blood sampling (week 0-week4 in a-b; week 4-week 8 in c-d). In all panels, dots represent individual samples.

## 4. Discussion

In this study we controlled the expression of autumn pre-migratory fattening in age-matched adult common quails and showed that Leucocyte Coping Capacity (LCC) peak responses varied markedly along the pre-migratory fattening process. We found that LCC was highest in the mid-fattening phase (week 4), but when quails achieved their peak fat load (week 8), LCC responses decreased to levels even lower than before the start of pre-migratory fattening (week 0). These relatively rapid, non-linear changes in LCC peak responses were somewhat surprising and only partly supported the general hypothesis that birds, as other migrating organisms, suppress immune system functioning before migration (Altizer et al., 2011; Eikenaar and Hegemann, 2016; Owen and Moore, 2006; Rogers et al., 2022).

The immunosuppression indicated by the low LCC response at the peak of pre-migratory fuelling is for example in line with a previous study on three species of thrushes showing that migrants had lower parameters of constitutive innate immune functions like leukocyte and lymphocyte counts compared to birds in breeding conditions (Owen and Moore, 2006). On the other hand, the marked increase in LCC peak levels we found four weeks after the beginning of pre-migratory fattening was not expected. It suggests that the immune system undergoes rapid remodelling as birds prepare for long-distance flights. Variation in immune responses over larger spans of the life cycle has been previously reported in birds. For example, a study in wild skylarks *Alauda arvensis* showed that some components of innate immune defence (hemolysis and hemagglutination) were highest during pre-migratory moulting, while concentrations of monocytes and basophils increased during autumn migration, and eosinophil concentration was highest during spring migration (Hegemann et al., 2012). However, the measurements of immune markers from the latter work were not the same across the study years and the birds’ age could not be estimated, which adds a further level of complexity for data interpretation (Buehler et al., 2009). On the other hand, a study of red knot *Calidris canutus* under controlled conditions using birds of similar age clearly showed changes in immune strategies across different stages of the annual cycle, with elevated indices of immunocosts during the period of body mass change (fattening and spring migration) compared to the periods when body mass was stable (Buehler et al., 2008). We propose that examinations over a shorter time scale like in the present study can be extremely useful to understand their potential link with behavioural and metabolic strategies characterising the distinct stages of the migratory period.

We found that changes in body composition along pre-migratory fattening were in part related to the changes in LCC peak responses. The Echo-MRI data confirmed that pre-migratory fuelling was mostly related to intense accumulation of fat stores and less to increase of lean mass, which would at least in part reflect increases in pectoral flight muscles (Boswell et al., 1993; Guglielmo, 2018; Price et al., 2011). We did find a significant increase in lean mass but this effect was limited to the first four weeks of the fattening process. We found a relationship between LCC peak and changes in body composition only at week 4 and this was more strongly related to changes in lean mass rather than fat mass. The birds showing the highest LCC peaks at week 4 kept a higher proportion of lean mass, and therefore also accumulated smaller fat stores. This result might be due to strategic shifts between metabolic remodeling (catabolic/anabolic) and keeping appropriate innate immune responses (Norris and Evans, 2000; Sheldon and Verhulst, 1996). Alternatively, birds that showed the highest LCC peaks may have delayed the accumulation of fat stores due to physiological constraints possibly associated to oxidative stress. A previous study in quails showed that the experimental control of the migratory state is associated with pronounced tissue-specific changes in oxidative status with pectoral muscle antioxidants (glutathione peroxidase) being positively related to increases in subcutaneous fat stores as the birds transitioned to a migratory state, and negatively related to losses in fat stores as the birds transitioned back to a non-migratory state (Marasco et al., 2021). Activated leucocytes produce free oxygen radicals in the frame of the so-called oxidative burst (McLaren et al., 2003) and reactive oxygen species play an important role in innate and specific immunity (Bogdan et al., 2000; Kohchi et al., 2009). Thus, decreasing LCC as well as overall number of leucocytes may be an important component of oxidative balance regulation protecting organism from oxidative damage and inflammation during energy challenges such as migration (Cywińska et al., 2010; Dick and Guglielmo, 2019; Nieman, 2000; Voigt et al., 2020). In future work, experimental manipulations involving changes in resources (for instance by limiting food availability) during migratory fuelling would be needed to test the prediction of physiological trade-offs with immune function. We also point out that studies using multiple markers/indices of both innate and acquired immunity would be important to obtain detailed information on seasonal remodelling of the immune system and its function in protecting individuals from disease and infectious risk during the energetically demanding phases of their life cycle.

## Acknowledgements

We thank the animal care team, especially Sabrinna Mali and Julia Kaiser, for their excellent support with animal husbandry, Roland Sasse for help with building the cages, Stefan Graf for technical assistance during the experiment, Thomas Paumann and Jonas Kahlen for technical support with the general maintenance of the husbandry facilities, Renate Hengsberger for her support with the formatting of the references, and Dr Rubina Mian for helpful discussions.

## Competing interests

Authors declaire no competing interests.

## Funding

The study was funded by FWF Der Wissenschaftsfonds Lise Meitner Fellowship (M2520-B29) to VM. MT was a scholarship holder funded by Polish National Agency for Academic Exchange: PPN/BEK/2020/1/00426. VM is most grateful to “NAWA PROM PROGRAMME” that supported her visit at the Department of Zoology (Poznań University of Life Sciences, Poland) during the drafting of the manuscript.

## Data availability

Data will be made publicly available upon acceptance of the manuscript.

## Author contributions (CRediT taxonomy)

Conceptualization: MT, VM

Methodology: ZZ, LF, NH, VM

Formal analysis: MT, VM

Investigation: MT, ZZ, IM, VM

Resources: LF, NH, GP, VM

Data curation: ZZ, MT, VM

Writing – original draft preparation: MT, ZZ, VM

Writing – review and editing: MT, LF, NH, IM, VM

Visualizazion: MT, VM

Supervision: VM

Project administration: VM

Funding acquisition: VM, LF

## Notes

### Competing Interest Statement

The authors have declared no competing interest.

